# Sex Differences in Auditory Brainstem Responses in the Hispid Pocket Mouse (*Chaetodipus hispidus*)

**DOI:** 10.1101/2024.12.13.628342

**Authors:** Luberson Joseph, Desi M. Joseph, Sarah Hobbs, Naleyshka Colon Rivera, Elizabeth A. McCullagh

## Abstract

The hispid pocket mouse (*C. hispidus*) is a solitary semi-fossorial rodent that has been the subject of various ecological and genetic studies. However, no previous studies have characterized its hearing ability, which is important for its survival and fitness. We collected auditory brainstem responses (ABRs) from *C. hispidus* and measured craniofacial and pinna morphological features to assess hearing ability and test differences in hearing thresholds, monaural and binaural ABR amplitudes and latencies between the sexes. ABR recordings revealed that similar to other small mammals, *C. hispidus* displayed the lowest threshold to sounds between 8-16 kHz, indicating best hearing across those frequencies. We found significant differences in auditory thresholds of the ABRs between the sexes, with females showing lower frequency hearing compared to males. However, no significant differences were detected in monaural and binaural ABR amplitudes and latencies between the sexes. We also found no significant differences in craniofacial and pinna dimensions between the sexes. These findings shed novel insights into the auditory systems across species and highlighted for the first time sex differences in auditory thresholds for this rodent species.

## INTRODUCTION

Hearing is a critical sensory modality and refers to the ability to perceive and transduce sound stimuli to the central auditory nervous system. For all taxa, this ability is vital for fitness and survival enabling foraging, predator escaping, prey and mate detection, navigation of habitats, and conspecific communication (Kidd et al. 1995; Grothe et al. 2010; Pfaff et al. 2015; Pleštilová et al. 2016; Potapova 2019; Scarpitti and Calede 2022). For spatial location and perception of sound sources, most taxa with external pinnae use two cues, interaural time differences (ITDs) and interaural level differences (ILDs). These cues are generated by timing or level differences between the two pinnae due to head size and pinna shape (Blauert 1997; Grothe et al. 2010). These cues are processed in the auditory brainstem, which translates ITD and ILD information into sound location. Which cues are available for sound localization processing of different taxa and their hearing ranges are critical to enhance our understanding of the variability of the auditory system among different groups of taxa.

The auditory brainstem response (ABR) is a useful technique to measure hearing sensitivity (i.e. hearing threshold, monaural / binaural amplitude and latency responses) (Bachmann and Hall 1998; Kim 2001; Zhou et al. 2006). In humans and other animal models, ABR recordings generate a characteristic waveform, with specific waves corresponding to the activation of ascending auditory areas within the peripheral and central auditory system (Popelar et al. 2008; Alvarado et al. 2012). ABRs capture evoked potentials generated from the auditory nerve to the midbrain, generally occurring within the initial 10 milliseconds following sound stimulation (Popelar et al. 2008). When performing ABRs on small mammals, four to five waves (reflecting synchronized neural responses) can be identified. In rodents, ABRs waves tend to vary in shape and amplitude across taxa, with the first wave often being the most prominent (Eggermont 2019). Thus, studies focusing on analyzing the ABRs wave patterns across taxa provides important insight into hearing ability of each species.

To date, limited studies have focused on comparing hearing ability between the sexes across rodent species. Indeed, males and females frequently exhibit differences in sensory processing owing to differences in developmental exposure to gonadal hormones and genetics (Arnold 2017; McCarthy et al. 2018). Female CBA/J and CBA/CaJ mice display lower ABR thresholds at 8 kHz and higher frequencies and showed faster decrease in hearing with aging compared to males (Henry 2004; Ohlemiller et al. 2010). Similarly, female Brattleboro and Long-Evans rats show lower ABR thresholds at frequencies 1, 4, 32, 42 kHz compared to male rats (Charlton et al. 2019). Sex-based differences in audiograms at high frequency in numerous mouse strains and rats, are influenced by factors such as aging, noise, hormones, and chemical exposure (Willott 2009; Lin et al. 2022). In addition, the degree of parental care and/or social grouping could also be factors leading to sex-differences in auditory perception among rodent species (Lin et al. 2022; New et al. 2024). For example, socially monogamous prairie voles (*Microtus ochrogaster*) exhibit sex differences in monaural amplitudes wave I and wave II, and monaural latency wave III and wave IV, although no differences in audiogram and binaural hearing ability (New et al. 2024). The Mongolian gerbil (*Meriones unguiculatus*) shows sex-differences in ABR threshold with aging, females exhibit lower auditory threshold compared to males at 16 kHz (Boettcher 2002). Therefore, investigating hearing ability of less social rodent species, especially those in which females provide parental care could provide novel insight into the relationship between parental care and hearing.

*Chaetodipus hispidus*, commonly known as the hispid pocket mouse is one of the most genetically divergent rodent species within the heteromyid genus *Chaetodipus* with four distinct subspecies and mitochondrial clades (Patton et al. 1981; Hafner et al. 2007; Andersen et al. 2012; Andersen and Light 2012). *C. hispidus* inhabits a large geographic area extending from North Dakota, through the grasslands of the Great Plains, and into the southwestern deserts of the United States (Paulson 1988; Geluso and Wright 2010). *C. hispidus* is commonly found in grazed and ungrazed grassland, shrubland, desert, and active and inactive cropland habitats with a diet consisting of a variety of seeds (i.e., forb, grass, succulent, etc.), insects, and green vegetation (Paulson 1988). *C. hispidus* are quadrupedal locomotors that live in underground borrow systems during the day and leave burrows only at night for mating and foraging (Kaufman et al. 2012).

Although previous work has focused on the genetics and ecology of *C. hispidus* (Paulson 1988; Andersen and Light 2012; Kaufman et al. 2012), to the best of our knowledge, no research has characterized the hearing range and ability to binaurally perceive sounds in this species. As an arid grassland specialist and solitary semi-fossorial species (Paulson 1988; Kaufman et al. 2012), like many species, hearing is of paramount importance for its survival and reproductive fitness. Previous studies have shown that hearing range of exclusive subterranean rodents is generally limited to low frequencies (between 0.5 – 1 kHz), with poor hearing and airborne sound sensitivity (Heffner and Heffner 1993; Kössl et al. 1996; Brückmann and Burda 1997). As a semi-fossorial rodent, middle ear structure of the hispid pocket mouse would favor best hearing thresholds of between 8 and 16 kHz similar with other semi-fossorial rodent like prairie voles (New et al. 2024).

In the current study, we used auditory brainstem responses (ABRs) to measure hearing sensitivity between sexes of the granivorous hispid pocket mouse, *C. hispidus* (Rodentia: Heteromyidae). We hypothesized that female hispid pocket mice will exhibit more sensitive hearing than males, characterized by lower auditory thresholds across frequencies, larger monaural amplitudes, and shorter latencies due to females being the primary responsible for parental care and therefore potentially more sensitive to sounds related to reproduction such as pups. We also hypothesized that there would be no differences in craniofacial dimension features or binaural-specific measures of latency and amplitude between the sexes.

## MATERIALS AND METHODS

### Ethics statement

All procedures used in this investigation were approved by the Oklahoma State University Institutional Animal Care and Use Committee (IACUC), were in accordance with the guidelines and recommendations of the American Society of Mammologists for the use of wild mammals in research (Sikes and the Animal Care and Use Committee of the American Society of Mammalogists 2016), and permission from an Oklahoma Department of Wildlife Conservation scientific collecting permit.

### Animals

Wild male and female hispid pocket mouse (*C. hispidus*) (N = 20, 10 males, 10 females) were live-trapped from two different locations in Oklahoma (Packsaddle Wildlife management areas, Arnet, OK (35^0^ 55’ 53.1” N, 99^0^ 44’ 24.5” W), and Selman Living Laboratory, Freedom, OK (36^0^ 42’ 46.9” N, 99^0^ 15’ 27.2” W) between May 2022 to August 2024 using aluminum Sherman (H.B Sherman Traps, Inc. Tallahassee, FL) non-folding traps (3” x 3” x 10”). All traps were baited with a mixture of creamy peanut butter and old-fashioned oats. Traps were generally positioned in vegetation or brush to minimize exposure to sun or rain and contained three cotton balls during the period from mid-December to mid-May of each year for warmth. The traps were left overnight and checked the following morning after approximately 12 hours. Upon capture, animals were transferred into a large unsealed plastic Ziploc bag for sex and species identification according to (Caire et al. 1989). Animals were next transferred into an Innovive (Animal Specialties and Provisions, LLC, Quakertown, PA) disposable plastic mouse cage (14.7” L x 9.2” W x 5.5” H) and transported to the laboratory at the Oklahoma State University for auditory brainstem response (ABR) recording.

### Auditory Brainstem Response (ABR) Recordings

Auditory brainstem responses were collected from both male and female *C. hispidus* to measure brainstem responses to sounds following similar techniques as previous publications (McCullagh et al. 2020; Chawla and McCullagh 2022; New et al. 2024). *C. hispidus* were initially anesthetized with a mixture of 60 mg/kg ketamine and 10 mg/kg xylazine and kept sedated for the duration of the procedure with a dosage of 25 mg/kg ketamine and 12 mg/kg xylazine. After being totally anesthetized, as indicated by lack of toe pinch reflex, animals were placed on a water-pump heating pad and kept at 37° C inside a small sound-attenuating chamber (Noise Barriers Lake Forest, IL, USA) lined with SONEX Tec Sound Proofing Wedges Foam in the interior (Illbruck Inc., Minneapolis, MN). ABR measurements were acquired by positioning three subcutaneous needle electrodes (Viasys Healthcare, Madison, WI, USA) under the skin of the sedated animals at the midline between the pinnae over the brainstem between (i.e., apex, active electrodes), the vertex (i.e., directly behind the apex on the nape, reference electrode), and in the back leg for ground. To record data from the inserted subcutaneous electrodes, we used a Tucker-Davis Technologies (TDT, Alachua, FL, USA), RA4LI head stage, a RA16PA preamplifier, and Multi I/O processor RZ5 attached to a PC with custom Python software.

The recorded data were processed using a second-order 50-3000 Hz filter and were averaged across 10-12 ms of recording time over 500 – 1000 repetitions. Tucker-Davis Technology Electrostatic Speakers (TDT EC-1) and Tucker-Davis Technology Electrostatic Speaker-Coupler Model (TDT MF-1) were used to present acoustic stimuli of 32 to 64 kHz and 1 to 24 kHz of frequencies/click stimuli respectively. Custom earphones were paired with an Etymotic ER-7C probe microphone (Etymotic Research Inc. Elk Grove, IL) for calibration of sound within the ear. Tone and click stimuli were presented to each animal with alternating polarities. Tone stimuli were 4 milliseconds in time with a 1 ms on-ramp and 1 ms off-ramp (2 ms ± 1 ms on/off ramps) while clicks were 0.1 milliseconds in duration. Acoustic stimuli were presented to the sedated animals at a 30 milliseconds interstimulus interval with a standard deviation of five milliseconds (Laumen et al. 2016) and were created at a sampling rate of 97656.25 Hz through a TDT RP2.1 real-time processor controlled by the custom Python program. Randomizing interstimulus intervals between each presentation has been shown to enhance the ABR waveform (Wang et al. 2020). Total ABR recording lasted approximately 120 minutes, after which animals were sacrificed using pentobarbital (30 mg/kg) followed by transcardial perfusion with phosphate-buffered saline (PBS) and paraformaldehyde (PFA).

### Auditory threshold, Monaural, and Binaural ABR Analyses

To determine the auditory threshold of animals, we used the method described by Brittan-Powell and Dooling 2004 for visual discrimination of auditory thresholds. This method consists of a trained researcher visually determined the lowest stimulus level per frequency that triggered an ABR response. Auditory thresholds were operationally described as the dB level halfway (5 dB) between the last visible ABR and the next lowest stimulus level. Auditory threshold stimuli consist of a 4 ms tone burst (2 ms ± 1 ms on/off ramp), of varying intensity and frequency. Click thresholds were also determined for each animal by decreasing click intensity in 5-10 dB SPL steps until the ABR waveforms totally disappeared, and similar to above, click threshold was defined as the intensity between absence of ABR waveforms and previously measured intensity (Figure 1).

**Figure 1:**
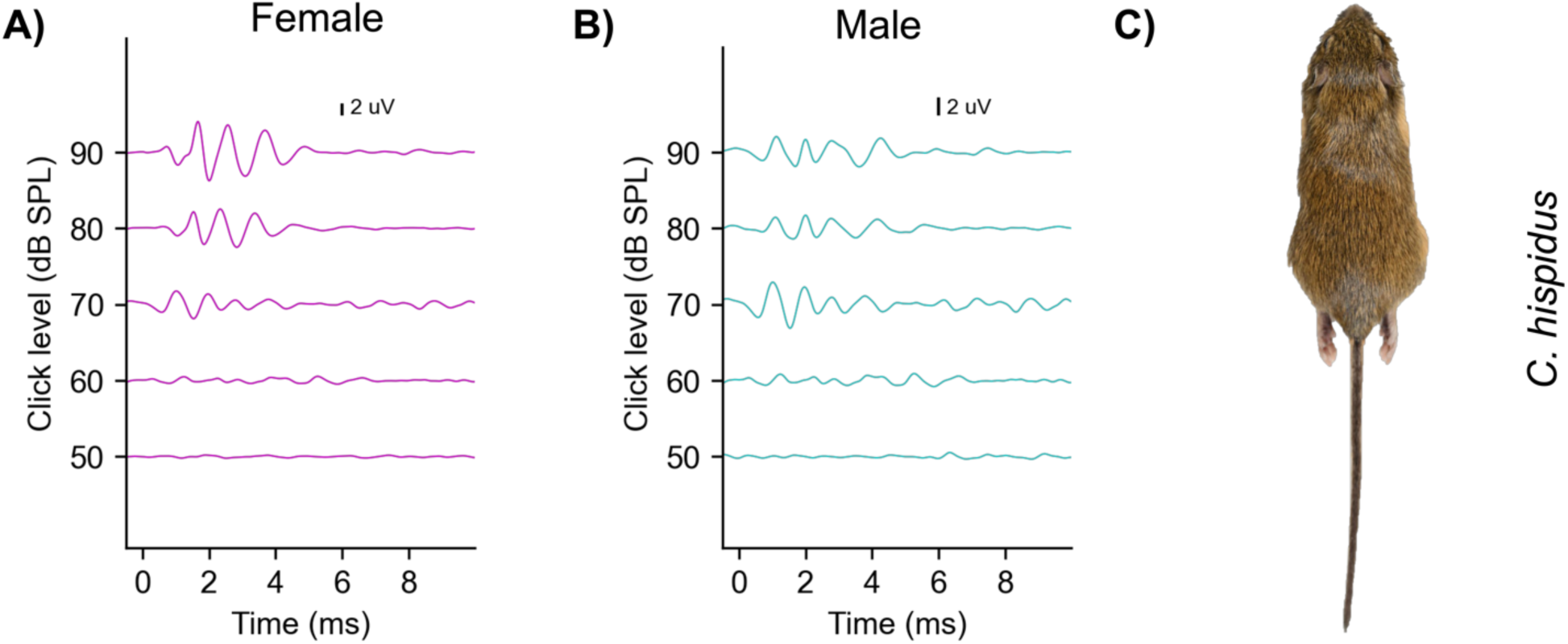
ABR traces to click stimuli for a representative female (magenta, left), and male (cyan, right) hispid pocket mouse. C is a photo of a female wild caught hispid pocket mouse (*C. hispidus*).

To generate monaural evoked potentials, we presented broadband click stimuli (i.e., 0.1 ms transient) at different intensities (60-90 dB SPL) to each pinna of the anesthetized animals and measured both peak latency (i.e., time to peak amplitude) and peak amplitude (i.e., the voltage from the peak to through) across the first four peaks of the ABR waveform.

Measurements of the monaural ABR latency and amplitude were calculated for each individual by calculating the average of the monaural latency and amplitude of wave I – IV from the auditory brainstem responses data acquired for sound presented at each pinna across intensities (60-90 dB SPL) (Zhou et al. 2006; New et al. 2024). Inter-peak latency between ABR peak I and peaks II, III, and IV were calculated as the difference in latency from the wave I to the other peak (II-IV) for right and left pinnae at intensities 60-, 70-, 80- and 90-dB SPL.

To evoke binaural auditory responses, we simultaneously presented broadband click stimuli at 90 dB SPL to both pinnae of the anesthetized animal. The binaural interaction component (hereafter called BIC) of the ABR was used as a measure of binaural sound processing and was measured by subtracting the sum of the two monaural auditory brainstem evoked responses from the binaural auditory brainstem response recordings. The BIC was described as the prominent negative peak (DN1) at wave four of the ABR following deduction of the summed monaural and binaural responses (Laumen et al. 2016; Benichoux et al. 2018). The BIC latency and amplitude were calculated using custom Python Software (New et al. 2024). To calculate BIC variation with ITD, animals were presented with click stimuli that ranged between– 2.0 to + 2.0 milliseconds ITD in 0.5 milliseconds steps. Corresponding BIC latencies and amplitudes were measured at each ITD. The amplitude of the BIC was determined as the peak relative to zero and baseline of the overall trace.

### Morphological Measurement Analysis

To compare morphological measurements between the sexes, seven standard external morphological measurements and body mass were taken for each individual with a six-inch Stainless Steel Electronic Vernier Caliper (DIGI-Science Accumatic digital Caliper Gyros Precision Tools Monsey, New York, USA) and a digital stainless Steel Electronic scale (Weighmax W-2809 90 LB X 0.1 OZ Durable Stainless Steel Digital Postal scale, Chino, California, USA), respectively. Standard morphological measurements included: body length (BL), tail length (TL), pinna length (PL), pinna width (PW), inter-pinna distance (IT), nose-to-pinna distance (NT), and effective pinna diameter (ED). Effective pinna diameter was calculated by taking the square root of the pinna width multiplied by the pinna length (Anbuhl et al. 2017).

Inter-pinna distance (mm) was measured as the distance between the two ear canals and the nose-to-pinna distance was calculated as the distance from the tip of the nose to the skull between the middle of the pinna. Pinna length was calculated as the distance between the basal notch to the tip (excluding hair), tail length and body length were measured as the distance between the sacrum to the caudal tip (excluding hair), and the distance from the tip of the nose to the caudal tip. All morphological dimensions data were recorded in thirteen juvenile (< 35 g; 6 males and 7 females,) and seven adult (≥ 35 g; 4 males and 3 females) *C. hispidus*, as estimated by their body mass to the nearest gram.

### Statistical Analysis

The audiogram, monaural peak amplitude and latency, inter-peak latency, and binaural auditory brainstem response data acquired between the sexes were compared statistically using linear mixed-effects models (LMMs). Audiogram, interpeak latency, monaural peak latency and amplitude, and BIC amplitude and latency across ITD were used as multivariate data with ITD, frequency, peak number, and sex as fixed effects and animal id as a random effect to account for multiple measurements within one animal. Estimated marginal means were used for pairwise comparisons of BIC amplitude and latency between both sexes (Russell 2018). Two-way Analysis of variance (ANOVA) was used to contrast differences in morphological measurements between the sexes. All analyses and figures were created in R Studio version 4.0.3 (R Core Team 2020), using the ggplot2 (Wickham 2016) and lme4 packages (Bates et al. 2014).

## RESULTS

As described above, we used tone bursts and click stimuli to compare the ABR threshold across frequencies (1-64 kHz), variation in monaural amplitudes and latencies, as well as variation in BIC amplitudes and latencies across ITDs (-2.0 to 2.0 ms) of ABR components between the sexes in wild caught hispid pocket mouse (C*. hispidus*). We also compare craniofacial and pinna morphology features including pinna length, pinna width, inter-pinna distances, nose-to-pinna distance, and pinna effective diameter, tail length, and body mass of hispid pocket mouse between the sexes.

### Auditory thresholds between sexes

ABR audiograms were constructed for females and males hispid pocket mouse to determine their sensitivity across frequencies. Both sexes exhibited lowest thresholds to tones in the 8 to 16 kHz region of the audiogram (Figure 2A), which is similar to other rodents of similar size (Lin et al. 2022; New et al. 2024). Linear mix-effect models revealed significant statistical difference between sexes across all frequencies tested (LMM: p-value = 0.021). When comparing specific frequency differences, female *C. hispidus* had significantly lower thresholds at 2 kHz (t-value = -2.403; p-value = 0.018), 4 kHz (t-value = -3.471; p-value = 0.0008), and 24 kHz (t-value = -2.461; p-value = 0.0143) compared to males; however, they showed similar hearing to males at all other frequencies tested (Table 1). We next investigated if there were sex differences in overall click thresholds. Similarly, we observed significant statistical differences in overall click threshold (p-value = 0.041) between the sexes, with females’ average click threshold displayed at 60 dB SPL, while that of males was 70 dB SPL (Figure 2B).

**Figure 2:**
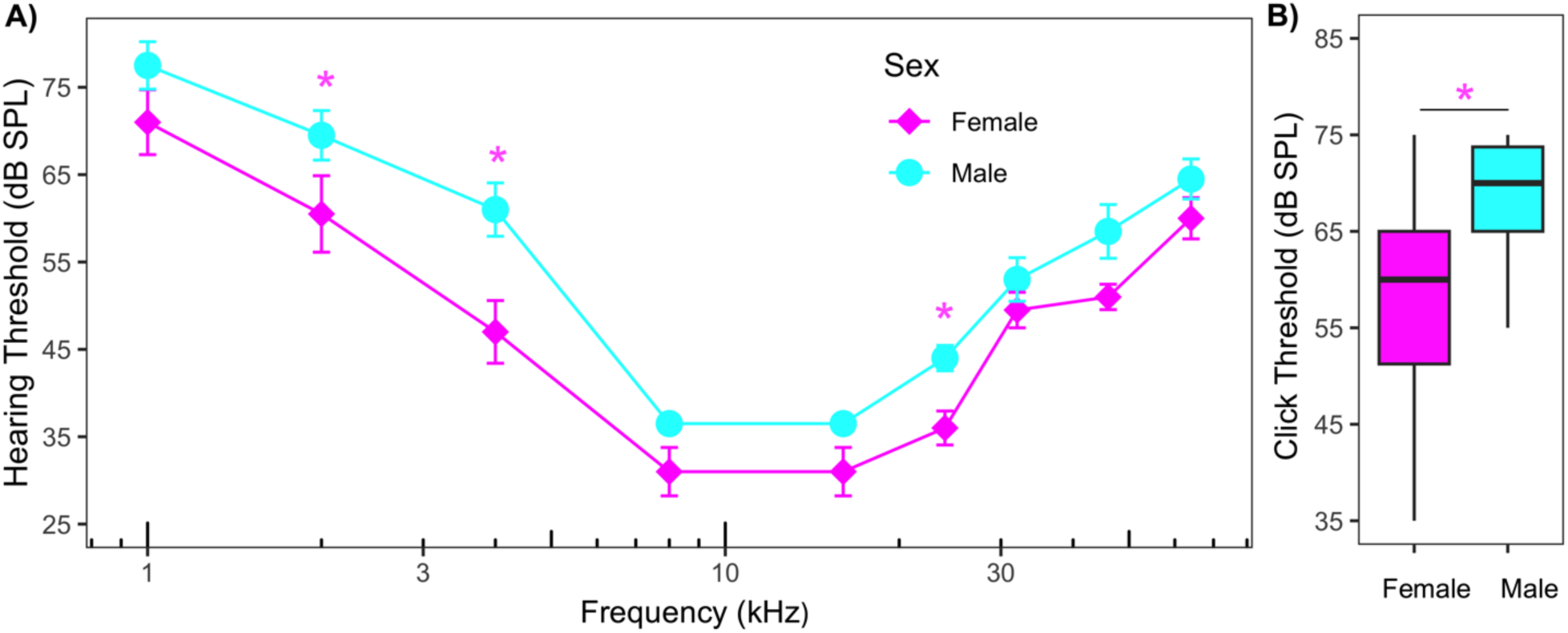
A) Audiogram of frequency sensitivity for male and female pocket mice, *C. hispidus* (N = 20, 10 males and 10 females) for tone stimulus ranging from 1 to 64 kHz. Significant main effects of frequency were detected, with significant differences between sexes at 2 kHz, 4 kHz, and 24 kHz ( *, all p-value < 0.05). Filled cyan circles represent males while filled magenta diamonds represent females. Figure B shows click thresholds for male and female pocket mouse. Significant differences were detected in click thresholds between sexes (*, p-value < 0.05).

**Table 1:** Linear mixed-effects model analysis and statistical significance testing comparing the ABR hearing thresholds of males (N = 10) and females (N = 10) *C. hispidus* for tone stimulus (mean ± Standard error, t-ratio, and p-value).

| Frequency (kHz) | Male | Female | t-ratio | p-value |
| --- | --- | --- | --- | --- |
| 1 | 76.5 ± 2.65 | 71 ± 2.65 | -1.469 | 0.146 |
| 2 | 69.5 ± 2.65 | 60.5 ± 2.65 | - 2.403 | 0.018 |
| 4 | 60 ± 2.65 | 47 ± 2.65 | - 3.471 | 0.0008 |
| 8 | 35.5 ± 2.65 | 31 ± 2.65 | - 1.202 | 0.233 |
| 16 | 35.5 ± 2.65 | 31 ± 2.65 | - 1.202 | 0.233 |
| 24 | 43 ± 2.65 | 36 ± 2.65 | - 2.461 | 0.0143 |
| 32 | 52 ± 2.65 | 49.5 ± 2.65 | - 0.668 | 0.506 |
| 46 | 57.5 ± 2.65 | 51 ± 2.65 | - 1.736 | 0.086 |
| 64 | 63.5 ± 2.65 | 60 ± 2.65 | - 0.935 | 0.353 |

### Monaural ABR waveform Amplitudes

We next measured the response of hispid pocket mouse to monaural transient click stimuli across intensities (60, 70, 80, 90 dB SPL) to determine if females and males share similar monaural hearing capability. We noticed that females tended towards slightly larger monaural peak amplitude than males at most intensities measured, although not statistically different (Figure 3; Table 2). For instance, at 90 dB SPL, the average amplitudes of waves I and IV were 2.98 and 1.53 μV, for female hispid pocket mouse while males were 1.52 and 1.21 μV, respectively (Figure 3). A linear mixed effect model revealed that both sexes displayed similar monaural wave amplitudes (I-IV) across intensities (LMM: p-value = 0.865), except in monaural wave II where females displayed larger monaural wave amplitude than males at 90 dB SPL (p-value = 0.038). However, we detected significant main effect of intensity on the monaural peak amplitude of waves I-IV for both sexes indicating that indeed amplitude increased with intensity across waves (LMM: p-value < 0.0001).

**Figure 3:**
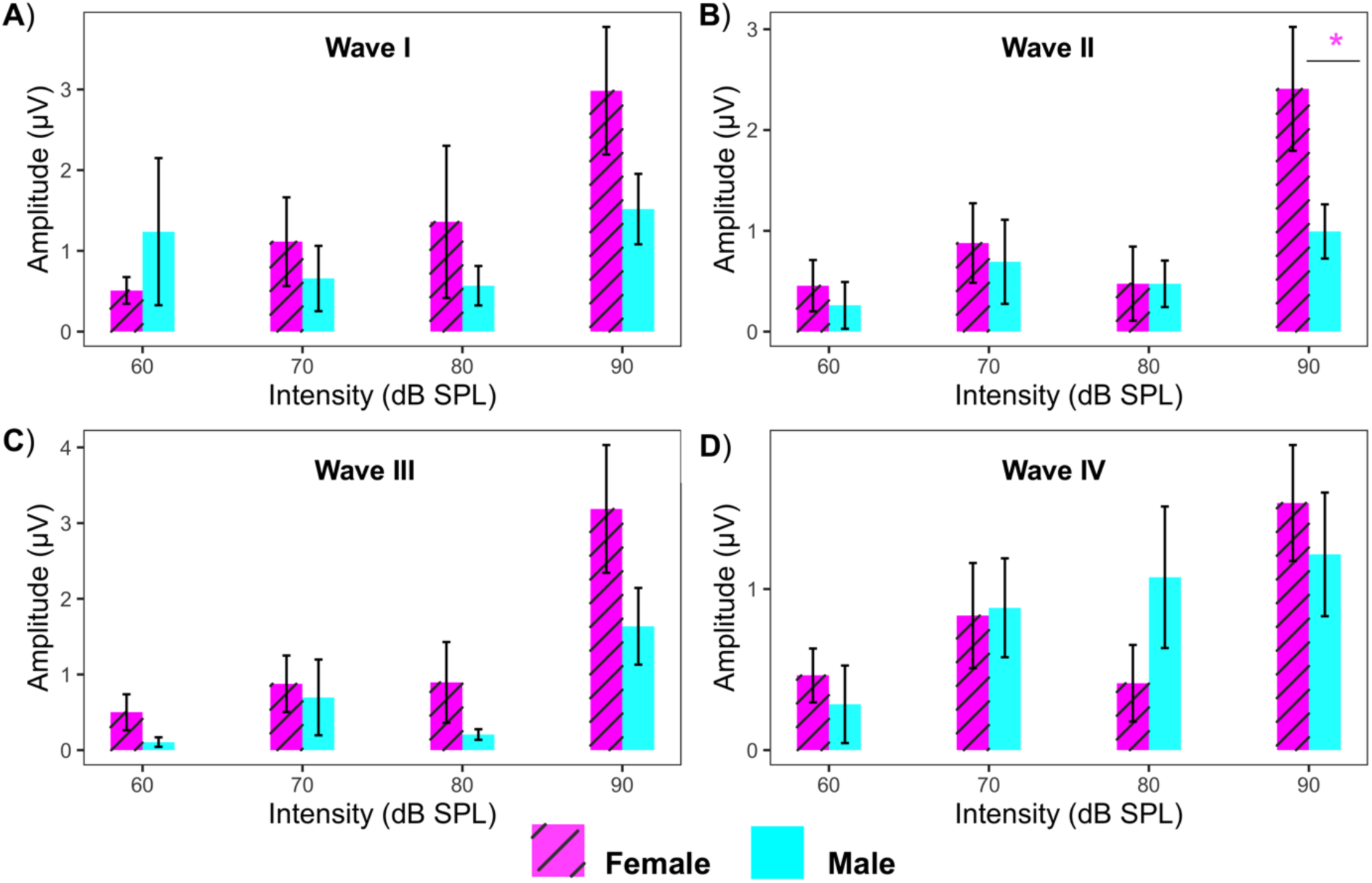
Average amplitude of ABR wave I-IV determined by clicks of different intensities between female and male hispid pocket mouse (*C. hispidus*) (Stripe magenta = Female (N=10), cyan = Male (N=10)). The vertical bars represent the standard error at each point. No significant difference of sex was detected on monaural ABR wave amplitude I, II, III and IV, but amplitude of waves did increase with intensity increases (p < 0.0001).

**Table 2:** Summary of monaural ABR amplitude of wave I-IV (mean ± Standard error, t-ratio, and p-value).

| Peak Amplitude ($\mu V$) | Male | Female | t-ratio | p-value |
| --- | --- | --- | --- | --- |
| I | $1.42 \pm 0.50$ | $1.67 \pm 0.52$ | 0.335 | 0.740 |
| II | $1.24 \pm 0.51$ | $1.38 \pm 0.52$ | 0.199 | 0.843 |
| III | $1.43 \pm 0.51$ | $1.68 \pm 0.53$ | 0.339 | 0.737 |
| IV | $1.35 \pm 0.50$ | $1.20 \pm 0.53$ | -0.208 | 0.837 |

### Absolute Latency

Absolute latencies were measured for waves I-IV across intensities ranging from 60 to 90 dB SPL for both female and male hispid pocket mouse. Like other small mammals, the hispid pocket mouse displayed a characteristic ABR, with four prominent peaks (I, II, III, and IV, see Figure 1). These peaks appeared at latencies comparable to those of other rodent species, occurring within the first 6 to 8 ms following the stimulation by the click stimuli (Benichoux et al. 2018a). The main effects of the linear mixed effects model did not show any significant effect of sex on ABR latency with increasing intensity (LMM: p-value = 0.948, Figure 4), however, there was a significant main effect of intensity on the absolute latency across the four ABR waves independent of sex suggesting that indeed latencies did increase with increases in intensity regardless of sex (LMM: p-value < 0.0001). However, there were no sex differences in peak latency as a function of intensity at any wave of the ABR (p-value > 0.05, Table 3).

**Figure 4:**
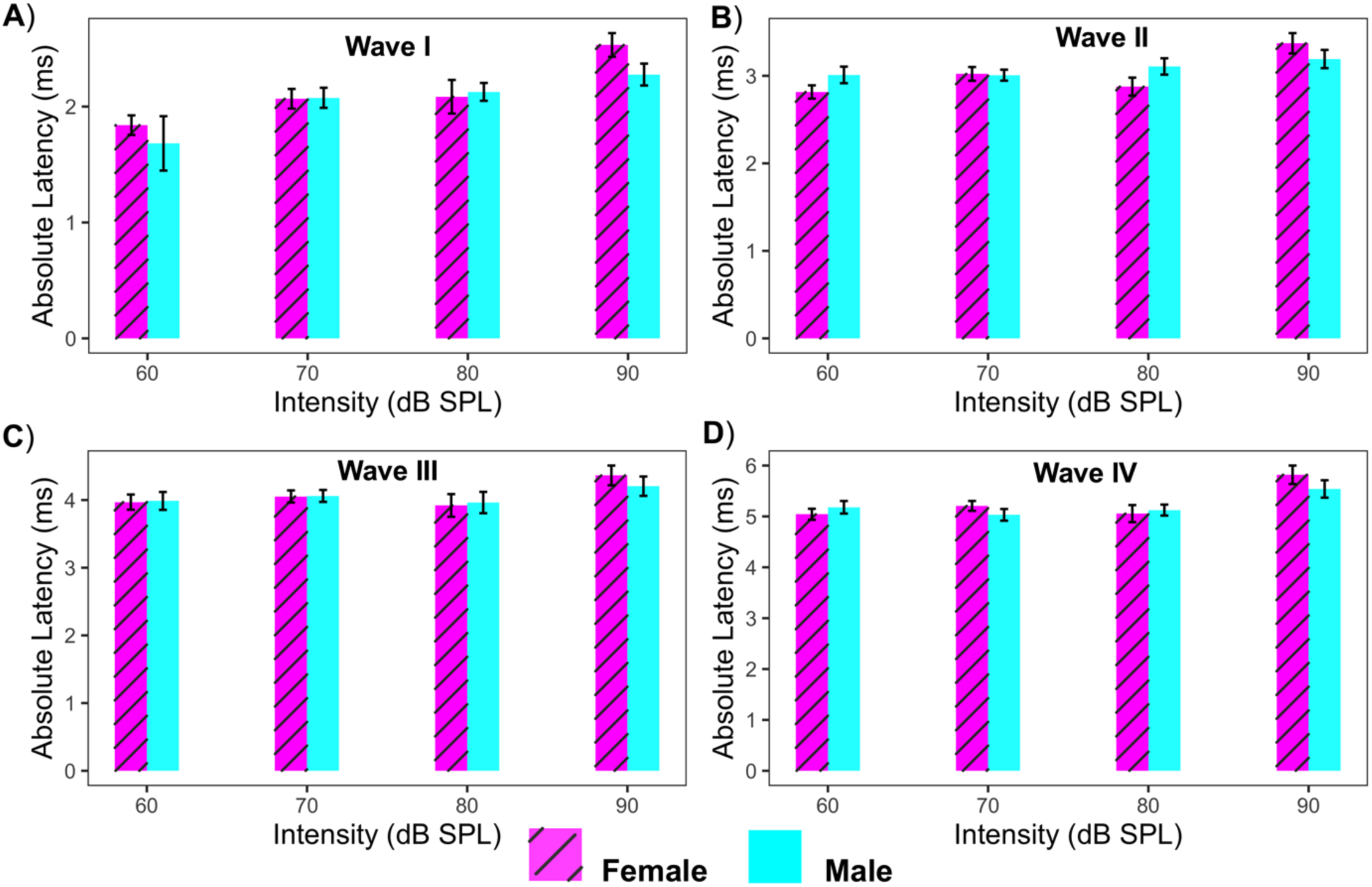
Average latency of ABR waves I-IV determined with clicks of different intensities between female and male hispid pocket mouse (*C. hispidus*) (stripe magenta = Female (N=10), cyan = Male (N=10)). The vertical bars represent the standard error at each point. No significant differences of sex were detected on monaural ABRs wave latency I, II, III, and IV, however there was a main effect of intensity on latency of the ABR waves (p-value < 0.0001).

**Table 3:**
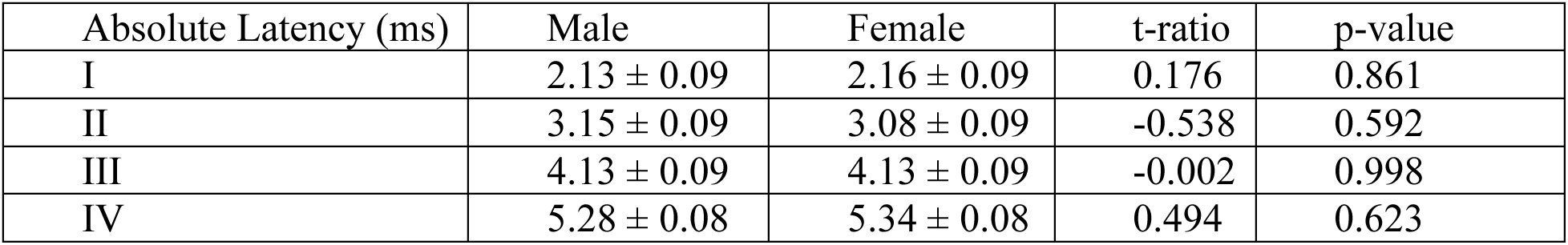
Summary of monaural ABR absolute latency of wave I-IV (mean ± Standard error, t-ratio, and p-value).

| Absolute Latency (ms) | Male | Female | t-ratio | p-value |
| --- | --- | --- | --- | --- |
| I | 2.13 ± 0.09 | 2.16 ± 0.09 | 0.176 | 0.861 |
| II | 3.15 ± 0.09 | 3.08 ± 0.09 | -0.538 | 0.592 |
| III | 4.13 ± 0.09 | 4.13 ± 0.09 | -0.002 | 0.998 |
| IV | 5.28 ± 0.08 | 5.34 ± 0.08 | 0.494 | 0.623 |

### Inter-peak Latency

Comparing sex effects on inter-peak latencies across intensities revealed no significant main effects of sex on inter-peak latency between male and female hispid pocket mice (LMM: p-value = 0.676, Figure 5), except at inter-peak latency I-II where females displayed shorter inter-peak latency than males at 80 dB SPL (p-value = 0.044). However, the linear mixed effects model showed a significant statistical effect of intensity on inter-peak latency for both sexes, indicating that indeed inter-peak latencies did decrease with increases in intensity regardless of sex (LMM: p-value = 0.032).

**Figure 5:**
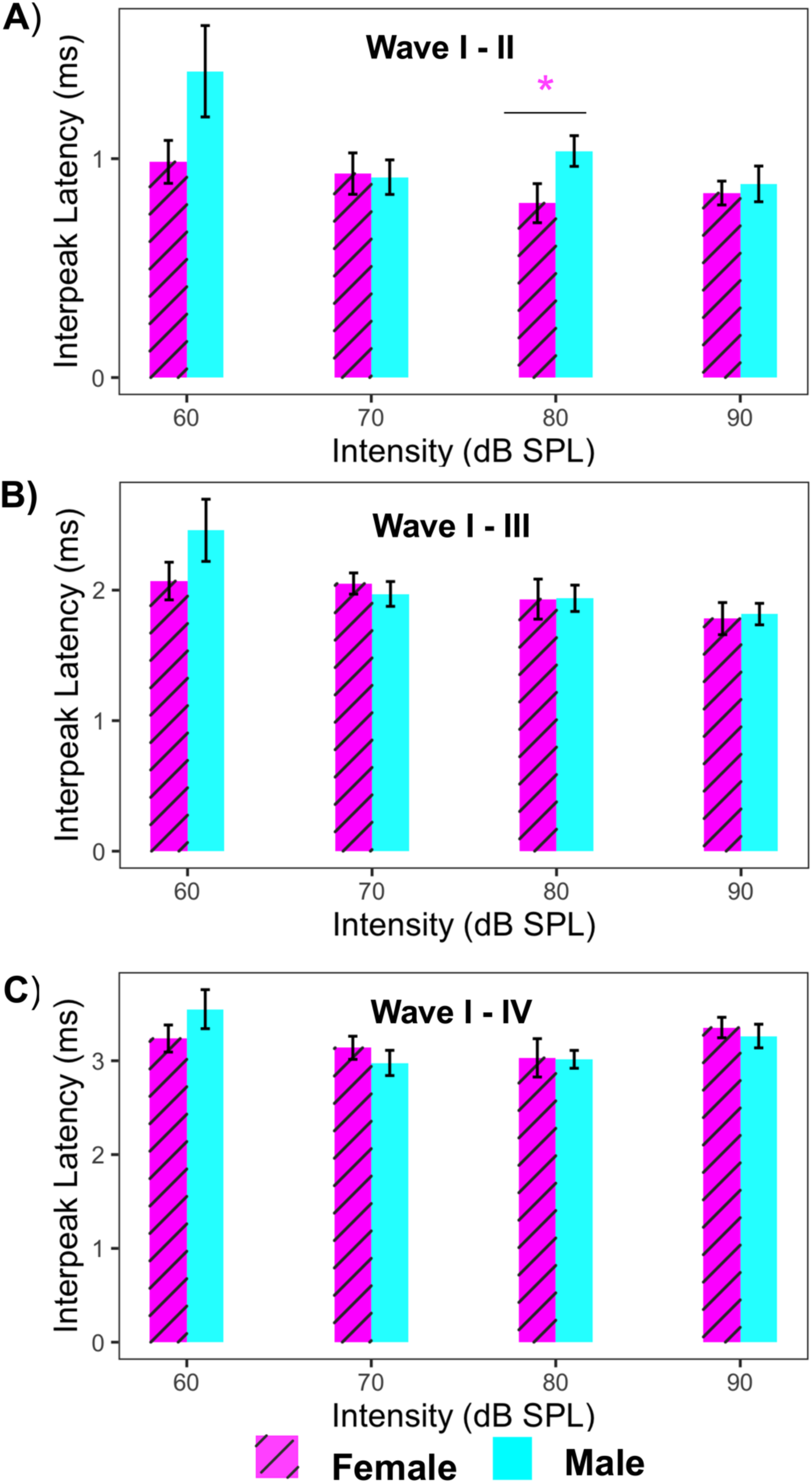
Average inter-peak latency of wave I-II, I-III, and I-IV determined with clicks of different intensities between female and male hispid pocket mouse (*C. hispidus*) (magenta stripe = Female (N=10), cyan = Male (N=10)). Error bars represent the standard error. No significant differences of sex were detected on monaural ABRs inter-peak latency I-II, I-III, and I-IV.

### Binaural ABR

Binaural processing in *C. hispidus’* was measured through binaural interaction component (BIC) amplitude and latency as it varies with ITD. The BIC is a measure of binaural hearing, and has been quantified across species (Laumen et al. 2016). Similar to previous findings, we observed that as ITD increases, BIC latency increases and amplitude decreases (Benichoux et al. 2018). Linear mixed effect model main effects did not reveal any overall significant statistical sex-specific effect of amplitude on the BIC as it varies across ITD ranging from -2 to +2 ms (Figure 6A; Table 4; LMM: p-value = 0.165). Similarly, we found no sex-specific differences in latency of the BIC as it varies across all ITDs (Figure 6B; Table 5; LMM: p-value = 0.177).

**Figure 6:**
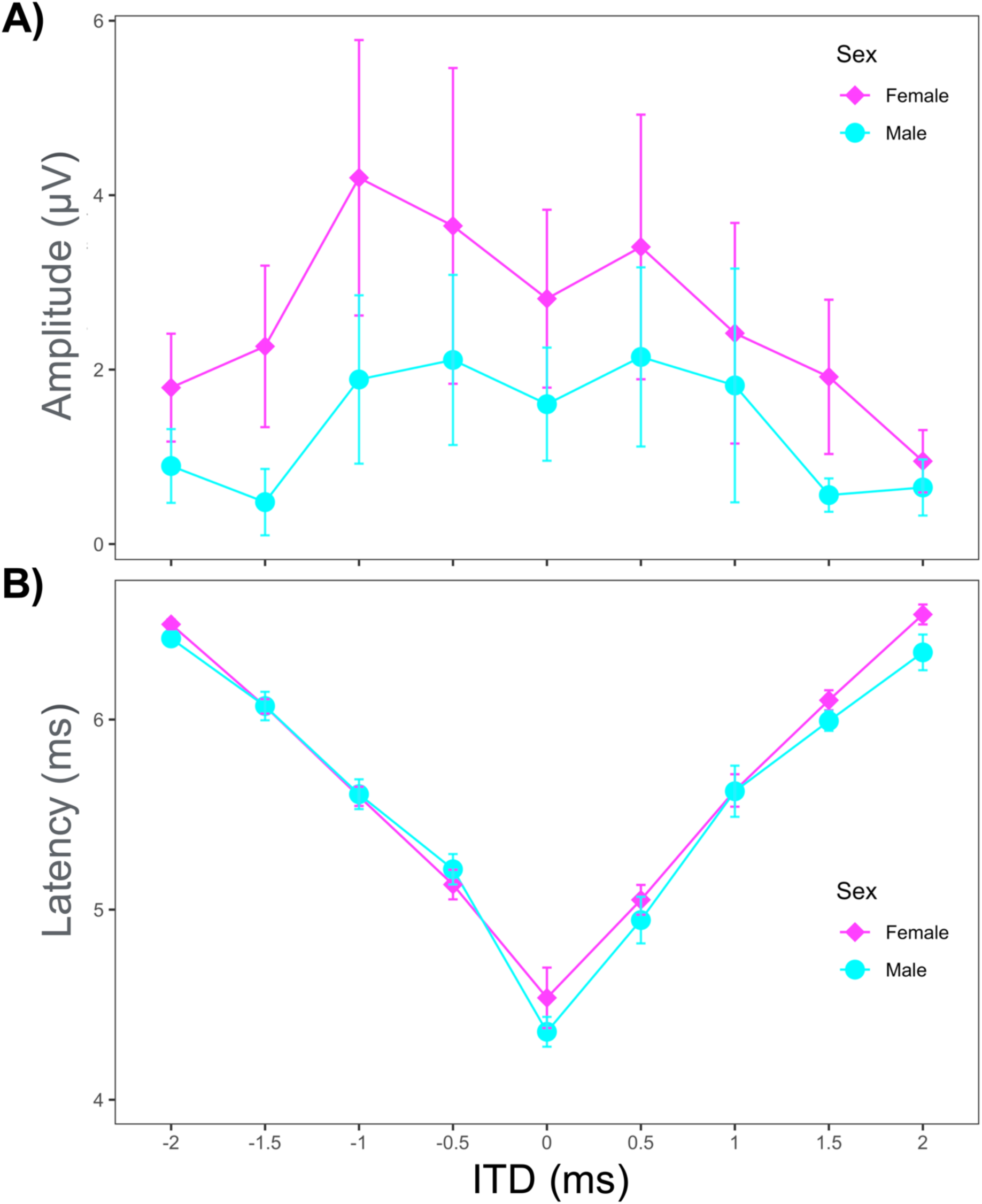
Binaural hearing comparison of female (filled magenta diamond) and male (filled cyan circle) *C. hispidus*. Binaural amplitude and latency of the BIC were measured for ITDs ranging from -2 to +2 ms in 0.5 ms steps. There were no sex differences in either overall BIC amplitude (Figure 6A) or BIC latency (Figure 6B) across ITDs measured. Error bars represent the standard error.

**Table 5:** Statistical measures of BIC amplitude as it varies with ITDs (mean ± standard error, t-ratio, and p-value).

| BIC amplitude (μV) | Male | Female | t-ratio | p-value |
| --- | --- | --- | --- | --- |
| -2.0 | $0.77 \pm 0.92$ | $1.79 \pm 0.99$ | 0.751 | 0.454 |
| -1.5 | $1.08 \pm 1.01$ | $2.66 \pm 0.99$ | 0.837 | 0.404 |
| -1.0 | $1.79 \pm 0.92$ | $3.93 \pm 1.04$ | 1.537 | 0.127 |
| -0.5 | $2.02 \pm 0.92$ | $3.68 \pm 1.07$ | 1.195 | 0.235 |
| 0.0 | $1.60 \pm 0.89$ | $2.81 \pm 0.99$ | 0.907 | 0.367 |
| 0.5 | $2.14 \pm 0.88$ | $3.41 \pm 0.99$ | 0.946 | 0.347 |
| 1.0 | $1.73 \pm 0.92$ | $2.41 \pm 1.04$ | 0.485 | 0.629 |
| 1.5 | $0.20 \pm 1.01$ | $1.92 \pm 0.99$ | 1.209 | 0.229 |
| 2.0 | $0.65 \pm 0.88$ | $0.95 \pm 0.99$ | 0.226 | 0.822 |

**Table 6:** Summary of BIC latency as it varies with ITDs (mean ± standard error, t-ratio, and p-value).

| BIC latency (ms) | Male | Female | t-ratio | p-value |
| --- | --- | --- | --- | --- |
| -2.0 | $6.41 \pm 0.08$ | $6.50 \pm 0.09$ | 0.724 | 0.470 |
| -1.5 | $6.09 \pm 0.09$ | $6.07 \pm 0.09$ | -0.126 | 0.901 |
| -1.0 | $5.61 \pm 0.08$ | $5.60 \pm 0.09$ | -0.073 | 0.941 |
| -0.5 | $5.21 \pm 0.08$ | $5.14 \pm 0.09$ | -0.565 | 0.573 |
| 0.0 | $4.36 \pm 0.08$ | $4.54 \pm 0.08$ | 1.506 | 0.134 |
| 0.5 | $4.95 \pm 0.07$ | $5.05 \pm 0.08$ | 0.897 | 0.371 |
| 1.0 | $5.62 \pm 0.08$ | $5.62 \pm 0.09$ | 0.011 | 0.991 |
| 1.5 | $5.97 \pm 0.09$ | $6.10 \pm 0.08$ | 1.008 | 0.315 |
| 2.0 | $6.35 \pm 0.08$ | $6.55 \pm 0.08$ | 1.681 | 0.095 |

### Morphological measurements

The external pinnae serve as the first component of the auditory system and are key in hearing variability across taxa. We measured numerous morphological features including pinna length, pinna width, effective pinna diameter, nose to pinna distance, inter pinna distance, body length, tail length, and body mass between the sexes. Two-way ANOVA showed no significant statistical differences in pinna measurements including pinna length (Figure 7B: F = 0.274, p-value = 0.607), pinna width (Figure 7C: F = 0.348, p-value = 0.563), and effective pinna diameter (Figure 7D: F = 0.538; p-value = 0.473) between the sexes. Similar to pinna dimensions, we detected no significant differences between sexes in craniofacial dimensions including inter-pinna distance (Figure 7E: F = 0.001; p = 0.97) and distance from the nose to the pinna (Figure 7F: F = 0.131; p = 0.721). Two-way ANOVA also detected no significant statistical sex-specific differences in body length (Figure 7G: F = 0.295; p = 0.593), tail length (Figure 7H: F = 0.747; p = 0.399), and body mass (Figure 7I: F = 0.285; p = 0.60).

**Figure 7:**
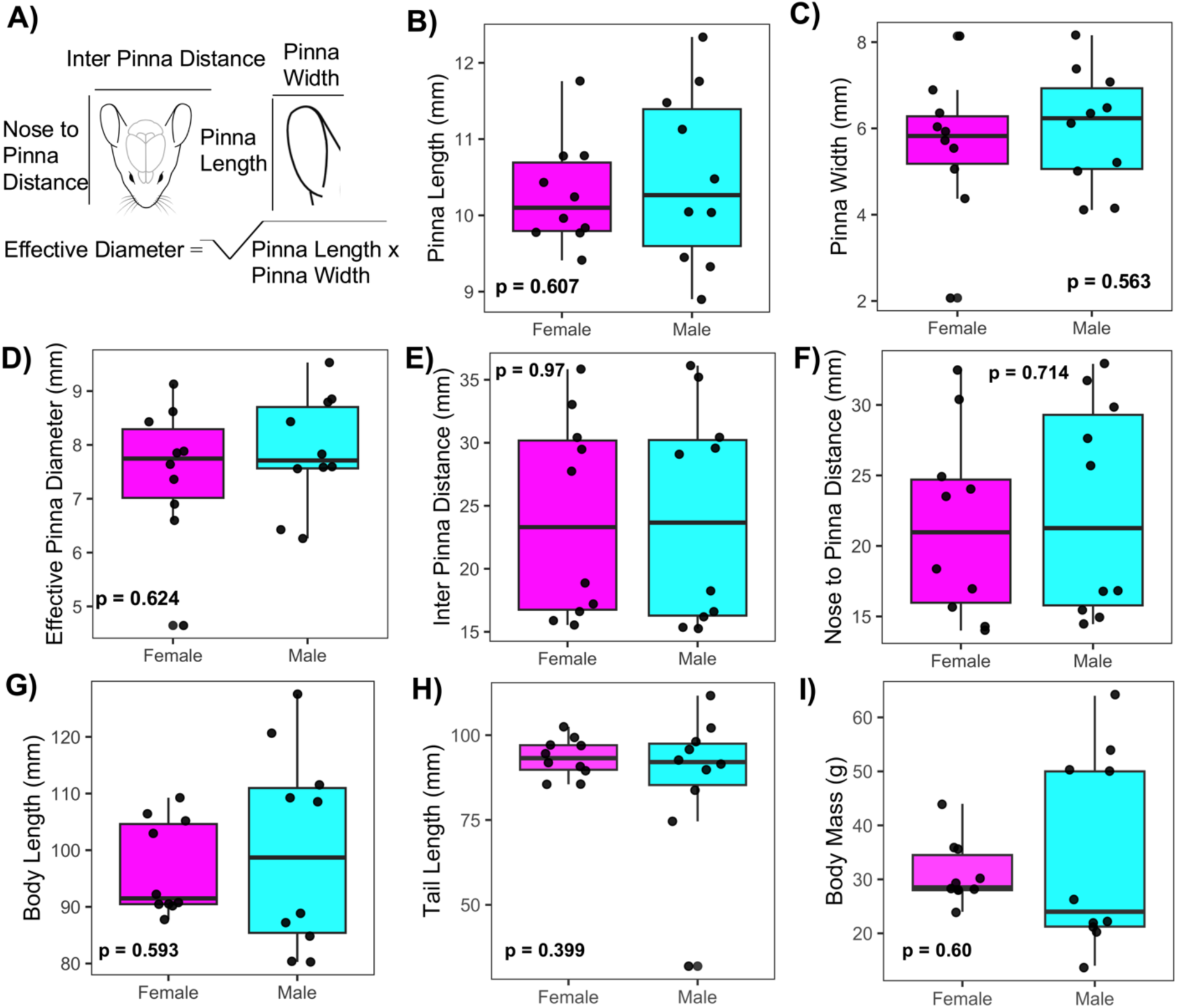
Morphological measurements in male and female *C. hispidus.* External pinna and head features were measured between sexes (magenta boxplot = female, cyan boxplot = male). No significant differences were detected in any morphological features measured between the sexes.

## DISCUSSION

This study represents the first comprehensive characterization of the auditory system of hispid pocket mouse (*C. hispidus*), including auditory thresholds, monaural and binaural hearing, and pinna and craniofacial morphology measurements between the sexes. While no significant sex differences were detected in monaural and binaural hearing ability or craniofacial and pinna morphology between male and female pocket mice, significant sex-specific differences were observed in monaural ABR and click thresholds, suggesting variation in auditory processing between males and females of the hispid pocket mouse, *C. hispidus* (Rodentia: Heteromyidae).

Pocket mice are of particular interest in phylogenetic and ecology studies due to their ability to inhabit arid environment, yet their auditory perception remains poorly understood. Hearing is crucial in rodents’ social communication, an area that has been largely overlooked in wild rodents including the hispid pocket mouse. The hearing thresholds observed in pocket mice aligns with findings from other semi-fossorial rodents (New et al. 2024) and perhaps with proposed auditory thresholds based on vocalization frequencies, though not documented in this species. However, this study is among the few to demonstrate sex differences in auditory thresholds in a semi-fossorial wild caught rodent species.

Significant main effects of sex in ABR thresholds were detected at selected low (2 and 4 kHz), and high frequencies (24 kHz), but were not consistent across all frequencies tested. It remains unclear whether these differences in ABR thresholds between the sexes occur at ecologically relevant frequencies. While 2 kHz and 4 kHz are not commonly associated with rodents ultrasonic vocalizations (USVs), 4 kHz falls within the range of rodents audible squeals, which are typically produced during distress and are thought to signal pain or discomfort (Borszcz 2006). The observed sex difference at 24 kHz ABR threshold is particularly significant, as this frequency is characteristic of ultrasonic vocalizations in rodents, which play an important role in social communication, including mating and parental care. Nonetheless, since this current study used pure tones rather than USVs, the alignment of pure tone frequencies with ultrasonic vocalization frequencies does not allow definitive conclusions regarding sex differences in USV detection between male and female hispid pocket mice. Therefore, future studies characterizing USV production, detection and behavioral responses between the sexes in this species will provide valuable insights into the role of auditory communication in the social and ecological behaviors of this species.

*C. hispidus* has been used as model organism in a number of studies due to its ability to inhabit arid ecosystems (Andersen and Light 2012; Andersen et al. 2012). However, data on hearing in this species has never been characterized. Our findings revealed that *C. hispidus*, despite being a semi-fossorial rodent, has similar hearing thresholds to predominantly surface-dwelling rodents including relatively high-frequency hearing (Dent et al. 2018; He et al. 2021). Comparative studies on hearing ability across different subspecies of the genus *Chaetodipus* are important to confirm if all subspecies share similar hearing as presented in this study. In addition, age has been reported to influence hearing sensitivity in many rodent species through increased auditory threshold and decreased high frequency hearing (Engle et al. 2013; Recanzone 2018). The subjects used in this experiment were captured from the wild, and therefore their precise ages were unknown, we were not able to assess the impact of age on auditory thresholds. Thus, future studies with captive-born animals would provide insight into age effects on threshold levels and high frequency hearing in hispid pocket mice.

Frequency thresholds of *C. hispidus* were similar to the thresholds described in other small mammals including the grasshopper mouse (*Onychomys leucogaster)*, and the Californian deer mouse (*Peromyscus californicus*) (Ralls 1967; Pasch et al. 2017; Capshaw et al. 2022).

Importantly, we should highlight that behavioral thresholds generally tend to be lower than those measured through ABR techniques and may be more variable between individuals. Previous studies comparing frequency thresholds between behavioral and ABR measures in human subjects revealed that behavioral thresholds are generally 10 -20 dB lower than those observed with ABR (Gorga et al. 1988; Stapells 2000). Nevertheless, ABRs allow for assessment of hearing in animals without behavioral training. Future behavioral experiments in the hispid pocket mouse (*C. hispidus*) would be useful to confirm ABR results in *C. hispidus*.

Most research on sex differences in small mammals’ ABRs highlight differential vulnerability to factors influencing hearing between the sexes (Lin et al. 2022). In addition, previous auditory studies of hearing show sex differences, typically manifested as changes in click thresholds or ABR thresholds, increased amplitude and decreased latency of wave II and IV in females (Debruyne et al. 1980; Jerger and Hall 1980; Edwards et al. 1983; Munro et al. 1997; Lin et al. 2022). Similarly, this study revealed significant differences in click threshold between the sexes in the hispid pocket mice, with female average click threshold was found to be 60 dB SPL, while that for males was 70 dB SPL. Our system of threshold detection is thought to be 10 to 20 dB SPL higher than measured due to the in-ear calibration, suggesting that the true hearing threshold for this species may be lower than described here. We detected no significant sex-specific differences in monaural amplitudes and latencies nor binaural amplitudes and latencies, as assessed by the binaural interaction component as it varies with ITD. While the present finding is consistent with previous studies demonstrating the shorter BIC latency and largest amplitude at zero ITD (Benichoux et al. 2018; Chawla and McCullagh 2022; New et al. 2024), our overall BIC-ITD association appear broader (Ferber et al. 2016; Benichoux et al. 2018), which may be attributed to differences in quantification techniques or other factors. Similar to previous findings, our data demonstrate a decrease in BIC amplitude with increasing ITD (Chawla and McCullagh 2022; New et al. 2024) and a corresponding increase in BIC latency with ITD. Individual craniofacial and pinna morphology should be considered when interpreting ABR results. Whereas none of the head morphology data were statistically different between the sexes, we conclude that head morphology measurements do not account for the monaural ABR differences between male and female pocket mice. Our hypothesis that females hispid pocket mice will exhibit more sensitive hearing than males, characterized by lower auditory thresholds across frequencies and larger monaural amplitude and shorter latency is partly supported.

Most small mammals rely on high-frequency hearing for sound location determination (Heffner and Heffner 2008). Factors such as functional interaural time and level differences (ITD and ILD) are useful for frequency-specific cues such as ITDs and ILDs and can be determined through morphological measurements (Masterton et al. 1969; Heffner and Heffner 2008). We calculated the functional interaural distance for our study species by summing the mean inter-pinna distance and pinna width then dividing by the speed of sound in air (331.29 m/s) to estimate the availability of ITD cues. With a mean inter-pinna distance of 15.8 mm and mean pinna width distance of 4.86 mm, we found that *C. hispidus* has a functional interaural time differences of about 63 μs. The relatively small pinna size compared to their head size may play additional roles in amplifying sound sources as the pinnae are moved, if they are motile. It is uncertain whether the hispid pocket mouse (*C. hispidus*) uses both ITD and ILD informations to pinpoint sound sources, as this species has not been tested in a behavioral setting. Although, we did not measure enough frequencies in the low-frequency range to determine the lowest limit of pocket mouse auditory range, thresholds of 71 -76 dB SPL detected at 1 kHz indicate poor low-frequency hearing, perhaps not sufficient to access ITD localization cues. These thresholds are higher than those reported for low-frequency rodents such as prairie voles (>50 dB SPL), and Mongolian gerbils (> 10 dB SPL), through direct comparisons between studies pose challenges due to differences in anesthesia effects and experimental setups (Smith and Mills 1991; New et al. 2024). Accordingly, the pocket mouse might not use ITDs to localize sound sources, but this will need to be confirmed behaviorally.

We found no significant sex differences in any morphological features measured including pinna length, pinna width, effective pinna diameter, nose-to-pinna distance, inter-pinna distance, body length, tail length, and body mass. Few studies have morphologically described *C. hispidus*. Two previous studies reported that adult of *C. hispidus* have a body weight of 16-80 g, body length of 80-120 mm, and tail length of 60-120 mm (Andersen and Light 2012; Kaufman et al. 2012), which were consistent with the current study. However, to our knowledge, no studies have reported craniofacial or pinna dimensions for this species. Accordingly, this study provides a detailed morphological characterization of the head and pinna of the hispid pocket mouse. In addition, head shape and size, and external pinnae size are important aspects that help small mammals hear and localize sound sources (Butler 1975; Grothe et al. 2010). Pinnae play a critical role in the removal of front-back confusion and detection of sound in the vertical plane (Heffner et al. 1996). Several rodent species display prominent secondary peaks for high-frequency hearing, and *C. hispidus* follows the same trend, with a secondary peak around 32 kHz. Accordingly, external pinnae size, shape, and directionality might couple with these secondary peaks as the smaller the pinna and head size are, generally the higher frequency of the species (Heffner and Heffner 2018). Thus, our second hypothesis that there would be no differences in craniofacial features or binaural-specific measures of latency and amplitude between the sexes is supported.

## CONCLUSION

We report for the first time sex differences in click threshold and auditory threshold across frequencies in the hispid pocket mouse (*C. hispidus*, Rodentia: Heteromyidae) using auditory brainstem responses (ABRs), craniofacial and pinna morphology. We show that the hispid pocket mouse (*C. hispidus*) exhibits a hearing range from 1-64 kHz, with optimal auditory sensitivity between 8 to 16 kHz. Importantly, we report few sex-specific differences in monaural or binaural hearing ability in this species, but significant differences in frequency sensitivity. Future studies incorporating behavioral and physiological assessments, particularly examining the effects of age impacts on auditory function, are essential to determine whether the observed sex differences in auditory thresholds, as well as monaural and binaural processing, are influenced by age-related factors in the hispid pocket mouse. If differences observed are not due to age effects, elucidating the mechanisms that underly differences in thresholds are an open area for subsequent experiments and may shed novel insights into hearing differences observed in other taxa.

## ACKNOWLEDGMENTS

Special thanks to Marcus Thibodeau and Marie Stone for housing and authorizing us to sample the packsaddle Wildlife Management Areas and the Selman Living Laboratory, respectively. We would like to thank the Payne County Audubon Society to LJ helped fund field sampling through the Helen Miller research grant.

## DATA AVAILABILITY

The data of the study will be made available upon request.

## AUTHOR CONTRIBUTIONS

LJ and EAM conceived and designed the experiments. LJ, DMJ, SH, and NCR trapped the rodents. LJ performed the experiments, analyzed the data, and wrote the first draft of the manuscript. All authors revised and edited the final manuscript.

## REFERENCES

Alvarado, J. C., V. Fuentes-Santamaría, T. Jareño-Flores, J. L. Blanco, and J. M. Juiz. 2012. Normal variations in the morphology of auditory brainstem response (ABR) waveforms: a study in wistar rats. Neuroscience Research 73:302–311.

Anbuhl, K. L., V. Benichoux, N. T. Greene, A. D. Brown, and D. J. Tollin. 2017. Development of the head, pinnae, and acoustical cues to sound location in a precocial species, the guinea pig ( Cavia porcellus). Hearing Research 356:35–50.

Andersen, J. J., and J. E. Light. 2012. Phylogeography and subspecies revision of the hispid pocket mouse, Chaetodipus hispidus (Rodentia: Heteromyidae). Journal of Mammalogy 93:1195–1215.

Andersen, J. J., M. A. Renshaw, and J. E. Light. 2012. Eight novel polymorphic microsatellites in the hispid pocket mouse (Chaetodipus hispidus) and cross-amplification in other Perognathinae species (Rodentia: Heteromyidae). Conservation Genetics Resources 4:1019–1021.

Arnold, A. P. 2017. A general theory of sexual differentiation. Journal of Neuroscience Research 95:291– 300.

Bachmann, K. R., and J. W. Hall. 1998. Pediatric Auditory Brainstem Response Assessment: The Cross-Check Principle Twenty Years Later. Seminars in Hearing 19:41–60.

Bates, D., M. Mächler, B. Bolker, and S. Walker. 2014. Fitting Linear Mixed-Effects Models using lme4.

Benichoux, V., A. Ferber, S. Hunt, E. Hughes, and D. Tollin. 2018a. Across Species “Natural Ablation” Reveals the Brainstem Source of a Noninvasive Biomarker of Binaural Hearing. The Journal of Neuroscience 38:8563–8573.

Benichoux, V., A. Ferber, S. Hunt, E. Hughes, and D. Tollin. 2018b. Across Species “Natural Ablation” Reveals the Brainstem Source of a Noninvasive Biomarker of Binaural Hearing. The Journal of Neuroscience 38:8563–8573.

Blauert, J. 1997. Spatial Hearing: The Psychophysics of Human Sound Localization. MIT Press.

Boettcher, F. A. 2002. Susceptibility to acoustic trauma in young and aged gerbils. The Journal of the Acoustical Society of America 112:2948–2955.

Borszcz, G. S. 2006. Contribution of the ventromedial hypothalamus to generation of the affective dimension of pain. Pain 123:155–168.

Brittan-Powell, E. F., and R. J. Dooling. 2004. Development of auditory sensitivity in budgerigars (Melopsittacus undulatus). The Journal of the Acoustical Society of America 115:3092–3102.

Brückmann, G., and H. Burda. 1997. Hearing in blind subterranean Zambian mole-rats (Cryptomys sp.): collective behavioural audiogram in a highly social rodent. Journal of Comparative Physiology A 181:83–88.

Butler, R. A. 1975. The Influence of the External and Middle Ear on Auditory Discriminations. Pp. 247–260 in Auditory System (W. D. Keidel & W. D. Neff, eds.). Springer Berlin Heidelberg, Berlin, Heidelberg.

Caire, W., J. D. Tyler, B. P. Glass, and M. A. Mares. 1989. Mammals of Oklahoma University of Oklahoma Press. Norman, Oklahoma 567.

Capshaw, G., S. Vicencio-Jimenez, L. A. Screven, K. Burke, M. M. Weinberg, and A. M. Lauer. 2022. Physiological Evidence for Delayed Age-related Hearing Loss in Two Long-lived Rodent Species (Peromyscus leucopus and P. californicus). Journal of the Association for Research in Otolaryngology 23:617–631.

Charlton, P. E., K. C. Schatz, K. Burke, M. J. Paul, and M. L. Dent. 2019. Sex differences in auditory brainstem response audiograms from vasopressin-deficient Brattleboro and wild-type Long-Evans rats. PLOS ONE 14:e0222096.

Chawla, A., and E. A. McCullagh. 2022a. Auditory Brain Stem Responses in the C57BL/6J Fragile X Syndrome-Knockout Mouse Model. Frontiers in Integrative Neuroscience 15.

Chawla, A., and E. A. McCullagh. 2022b. Auditory Brain Stem Responses in the C57BL/6J Fragile X Syndrome-Knockout Mouse Model. Frontiers in Integrative Neuroscience 15.

Debruyne, F., G. Hombergen, and M. Hoekstra. 1980. [Normal values in brain stem electric response audiometry (BERA). Acta oto-rhino-laryngologica Belgica 34:238–245.

Dent, M. L., L. A. Screven, and A. Kobrina. 2018. Hearing in Rodents. Pp. 71–105 in Rodent Bioacoustics (M. L. Dent, R. R. Fay & A. N. Popper, eds.). Springer International Publishing, Cham.

Edwards, R. M., N. K. Squires, J. S. Buchwald, and P. E. Tanguay. 1983. Central Transmission Time Differences in the Auditory Brainstem Response as a Function of Sex, Age, and Ear of Stimulation. International Journal of Neuroscience 18:59–66.

Eggermont, J. J. 2019. Chapter 30 - Auditory brainstem response. Pp. 451–464 in Handbook of Clinical Neurology (K. H. Levin & P. Chauvel, eds.). Elsevier.

Engle, J. R., S. Tinling, and G. H. Recanzone. 2013. Age-Related Hearing Loss in Rhesus Monkeys Is Correlated with Cochlear Histopathologies. PLoS ONE 8:e55092.

Ferber, A. T., V. Benichoux, and D. J. Tollin. 2016. Test-Retest Reliability of the Binaural Interaction Component of the Auditory Brainstem Response. Ear and hearing 37:e291–e301.

Fetoni, A. R., P. M. Picciotti, G. Paludetti, and D. Troiani. 2011. Pathogenesis of presbycusis in animal models: A review. Experimental Gerontology 46:413–425.

Geluso, K., and G. D. Wright. 2010. Hispid Pocket Mouse ( Chaetodipus hispidus) in East-Central and Northeastern Nebraska. Western North American Naturalist 70:126–129.

Gorga, M. P., J. R. Kaminski, K. A. Beauchaine, and W. Jesteadt. 1988. Auditory Brainstem Responses to Tone Bursts in Normally Hearing Subjects. Journal of Speech, Language, and Hearing Research 31:87–97.

Grothe, B., M. Pecka, and D. McAlpine. 2010a. Mechanisms of Sound Localization in Mammals. Physiological Reviews 90:983–1012.

Grothe, B., M. Pecka, and D. McAlpine. 2010b. Mechanisms of sound localization in mammals. Physiological Reviews 90:983–1012.

Hafner, J. C. et al. 2007. Basal Clades and Molecular Systematics of Heteromyid Rodents. Journal of Mammalogy 88:1129–1145.

He, K. et al. 2021. Echolocation in soft-furred tree mice. Science 372:eaay1513.

Heffner, H. E., and R. S. Heffner. 2008. High-Frequency Hearing. Pp. 55–60 in The Senses: A Comprehensive Reference. Elsevier.

Heffner, H. E., and R. S. Heffner. 2018. The evolution of mammalian hearing. AIP Conference Proceedings 1965:130001.

Heffner, R. S., and H. E. Heffner. 1993. Degenerate hearing and sound localization in naked mole rats (Heterocephalus glaber), with an overview of central auditory structures. Journal of Comparative Neurology 331:418–433.

Heffner, R. S., G. Koay, and H. E. Heffner. 1996. Sound localization in chinchillas III: Effect of pinna removal. Hearing Research 99:13–21.

Henry, K. R. 2004. Males lose hearing earlier in mouse models of late-onset age-related hearing loss; females lose hearing earlier in mouse models of early-onset hearing loss. Hearing Research 190:141–148.

Jerger, J., and J. Hall. 1980. Effects of Age and Sex on Auditory Brainstem Response. Archives of Otolaryngology 106:387–391.

Kaufman, G. A., D. M. Kaufman, and D. W. Kaufman. 2012. Hispid Pocket Mice in Tallgrass Prairie: Abundance, Seasonal Activity, Habitat Association, and Individual Attributes. Western North American Naturalist 72:377–392.

Kidd, G., C. R. Mason, and T. L. Rohtla. 1995. Binaural advantage for sound pattern identification. The Journal of the Acoustical Society of America 98:1977–1986.

Kim, K. P., Xiao-Ming Sun, Duck O. 2001. Noninvasive Assessment of Auditory Function in Mice: Auditory Brainstem Response and Distortion Product Otoacoustic Emissions. Handbook of Mouse Auditory Research. CRC Press.

Kössl, M., G. Frank, H. Burda, and M. Müller. 1996. Acoustic distortion products from the cochlea of the blind African mole rat, Cryptomys spec. Journal of Comparative Physiology A 178:427–434.

Laumen, G., A. T. Ferber, G. M. Klump, and D. J. Tollin. 2016. The Physiological Basis and Clinical Use of the Binaural Interaction Component of the Auditory Brainstem Response. Ear & Hearing 37:e276–e290.

Lin, N., S. Urata, R. Cook, and T. Makishima. 2022. Sex differences in the auditory functions of rodents. Hearing Research 419:108271.

Masterton, B., H. Heffner, and R. Ravizza. 1969. The Evolution of Human Hearing. The Journal of the Acoustical Society of America 45:966–985.

McCarthy, M. M., K. Herold, and S. L. Stockman. 2018. Fast, furious and enduring: Sensitive versus critical periods in sexual differentiation of the brain. Physiology & Behavior 187:13–19.

McCullagh, E. A., S. Poleg, N. T. Greene, M. M. Huntsman, D. J. Tollin, and A. Klug. 2020. Characterization of Auditory and Binaural Spatial Hearing in a Fragile X Syndrome Mouse Model. eneuro 7:ENEURO.0300-19.2019.

Munro, K. J., J. N. Shiu, and C. L. Cox. 1997. The Effect of Head Size on the Auditory Brainstem Response for Two Breeds of Dog. British Journal of Audiology 31:309–314.

New, E. M. et al. 2024a. Hearing ability of prairie voles (Microtus ochrogaster). The Journal of the Acoustical Society of America 155:555–567.

New, E. M. et al. 2024b. Hearing ability of prairie voles (Microtus ochrogaster). The Journal of the Acoustical Society of America 155:555–567.

New, E. M. et al. 2024c. Hearing ability of prairie voles (Microtus ochrogaster). The Journal of the Acoustical Society of America 155:555–567.

Ohlemiller, K. K., A. R. Dahl, and P. M. Gagnon. 2010. Divergent Aging Characteristics in CBA/J and CBA/CaJ Mouse Cochleae. Journal of the Association for Research in Otolaryngology 11:605– 623.

Pasch, B., I. T. Tokuda, and T. Riede. 2017. Grasshopper mice employ distinct vocal production mechanisms in different social contexts. Proceedings of the Royal Society B: Biological Sciences 284:20171158.

Patton, J. L., S. W. Sherwood, and S. Y. Yang. 1981. Biochemical systematics of chaetodipine pocket mice, genus Perognathus. Journal of Mammalogy 62:477–492.

Paulson, D. D. 1988. Chaetodipus hispidus. Mammalian Species no. 320 (1–4). American Society of Mammalogists New York.

Pfaff, C., T. Martin, and I. Ruf. 2015. Bony labyrinth morphometry indicates locomotor adaptations in the squirrel-related clade (Rodentia, Mammalia). Proceedings of the Royal Society B: Biological Sciences 282:20150744.

Pleštilová, L., E. Hrouzková, H. Burda, and R. Šumbera. 2016. Does the morphology of the ear of the Chinese bamboo rat (Rhizomys sinensis) show “Subterranean” characteristics? Journal of Morphology 277:575–584.

Popelar, J., J. Grecova, N. Rybalko, and J. Syka. 2008. Comparison of noise-induced changes of auditory brainstem and middle latency response amplitudes in rats. Hearing Research 245:82–91.

Potapova, E. G. 2019. Morphological Specificity of the Auditory Capsule of Sciurid (Sciuridae, Rodentia). Biology Bulletin 46:730–743.

Ralls, K. 1967. Auditory sensitivity in mice: Peromyscus and Mus musculus. Animal Behaviour 15:123– 128.

Recanzone, G. 2018. The effects of aging on auditory cortical function. Hearing Research 366:99–105.

Russell, L. 2018. Emmeans: estimated marginal means, aka least-squares means. R package version 1.

Scarpitti, E. A., and J. J. M. Calede. 2022. Ecological correlates of the morphology of the auditory bulla in rodents: Application to the fossil record. Journal of Anatomy 240:647–668.

Sikes, R. S. and the Animal Care and Use Committee of the American Society of Mammalogists. 2016. 2016 Guidelines of the American Society of Mammalogists for the use of wild mammals in research and education. Journal of Mammalogy 97:663–688.

Smith, D. I., and J. H. Mills. 1991. Low-frequency component of the gerbil brainstem response: Response characteristics and anesthesia effects. Hearing Research 54:1–10.

Stapells, D. R. 2000. Threshold estimation by the tone-evoked auditory brainstem response: a literature meta-analysis. Journal of Speech Language Pathology and Audiology 24:74–83.

Wang, X. et al. 2020. The Effects of Random Stimulation Rate on Measurements of Auditory Brainstem Response. Frontiers in Human Neuroscience 14.

Wickham, H. 2016. Programming with ggplot2. Pp. 241–253 in ggplot2: Elegant Graphics for Data Analysis (H. Wickham, ed.). Springer International Publishing, Cham.

Willott, J. F. 2009. Effects of sex, gonadal hormones, and augmented acoustic environments on sensorineural hearing loss and the central auditory system: Insights from research on C57BL/6J mice. Hearing Research 252:89–99.

Zhou, X., P. H.-S. Jen, K. L. Seburn, W. N. Frankel, and Q. Y. Zheng. 2006a. Auditory brainstem responses in 10 inbred strains of mice. Brain Research 1091:16–26.

Zhou, X., P. H.-S. Jen, K. L. Seburn, W. N. Frankel, and Q. Y. Zheng. 2006b. Auditory brainstem responses in 10 inbred strains of mice. Brain Research 1091:16–26.

